# Narwhal (*Monodon monoceros*) echolocation click rates to support cue counting passive acoustic density estimation

**DOI:** 10.1101/2023.05.23.541771

**Authors:** Carolina S. Marques, Diana A. Marques, Susanna B. Blackwell, Mads Peter Heide-Jørgensen, Chloe E. Malinka, Tiago A. Marques

## Abstract

Estimating animal abundance is fundamental for effective management and conservation. It is increasingly done by combining passive acoustics with knowledge about rates at which animals produce cues (cue rates). Narwhals (*Monodon monoceros*) are elusive marine mammals for which passive acoustic density estimation might be plausible, but for which cue rates are lacking. Clicking rates in narwhals were investigated using a pre-existing dataset from sound and movement tag records collected in August 2013-2016 and 2019 in East Greenland. Clicking rates were quantified for *∼*1200 one-second-long systematic random samples from 8 different whales. Generalized Additive Models were used to model (1) the probability of being in a clicking state versus depth; and (2) the clicking rate while in a clicking state, versus time and depth. The probability of being in a clicking state increased with depth, reaching *∼*1.0 at *∼*500 meters, while the number of clicks per second (while in a clicking state) increased with depth. The mean cue production rate, weighted by tag duration, was 1.28 clicks per second (se= 0.13, CV= 0.10). This first cue rate for narwhals may be used for cue counting density estimation, but care should be taken if applying it to other geographical areas or seasons.

## I. INTRODUCTION

Cetaceans are largely dependent on sound for interacting with their environment. They use different vocalizations to forage, communicate, escape from danger, orientate and navigate (Weilgart, 2017). Therefore, their sounds (e.g., echolocation clicks, burst pulses, whistles, etc.), which we will refer generically as cues, might reflect their behavioural state. Information on acoustic cues and the rates at which they are emitted can improve estimates of population density, as derived from acoustic data. Density estimates over time allow the detection of population trends, anthropogenic effects on the population, and the impacts of oceanographic and climate changes (Marques *et al*., 2013). For some species, surveying animals acoustically might prove more efficient than traditional visual methods. Cetacean sounds can be used to estimate their density using Passive Acoustic Monitoring (PAM) (e.g. Blackwell *et al*., 2018; Harris *et al*., 2018; Marques *et al*., 2009; Nowacek *et al*., 2016; Warren *et al*., 2017). A commonly used approach to do so is cue counting, where a density of cues (sounds of interest) is converted into a density of animals. To do so, an estimate of the average number of cues produced per animal per unit time - the cue rate is required.

Animal-borne tags are often used to collect relevant information about animal behaviour and physiology. Biologging tags can record information from a few hours to several months, providing a unique window of observation into the lives of otherwise elusive animals. These tags can sometimes also record animal movement, location, and/or environmental covariates around the tagged individual (Burgess *et al*., 1998; Johnson and Tyack, 2003). Many of these tags also record acoustic data, which makes them prime tools for collecting data to estimate relevant acoustic cue rates to inform passive acoustic density estimation exercises (Marques *et al*., 2013).

Several factors might affect PAM density estimates, including low population densities combined with wide variation in cue activity, or low cue production rates. These variabilities will lower the odds of reliably and consistently detecting cues on recorders, and hence spatiotemporal variability in detections will be high and sample sizes low (Blackwell *et al*., 2018; Marques *et al*., 2013; Scheidat *et al*., 2019). A reliable and adequate mean cue rate estimate, valid for the time and place a survey takes place and required for an accurate density estimate, is hard to obtain. The cue rate might be dependent on individual characteristics, like their reproductive status, age, sex, etc (Thomas and Marques, 2012), or might be seasonally and environmentally driven (Ladegaard *et al*., 2021). Therefore, for a reliable density estimation using PAM, above and beyond understanding a given species’ bio-sonar properties, such as the beam-width and frequency composition of the sounds used for cue counting (Macaulay *et al*., 2020), it is essential to have good knowledge of the species’ acoustic ecology and behaviour.

Narwhals (*Monodon monoceros*) are toothed whales with many unknowns regarding their acoustic ecology, including an undefined acoustic cue rate. Over time, narwhal sub- populations have slowly decreased in genetic diversity, and if this tendency continues, the survival of the species will be threatened (Westbury *et al*., 2019). Narwhals are particularly sensitive to climate change due to their constrained distribution in the Arctic and marked loyalty to their pack-ice habitat, which plays an important role during winter feeding (Laidre *et al*., 2008; Shuert *et al*., 2022; Williams *et al*., 2011). Climate change is leading to rises in sea temperature and ice melting, which has consequences on narwhal habitat availability. This warming has been predicted to lead to the disappearance of summer ice in the Arctic Sea during our lifetime (Stocker *et al*., 2014). This is especially of concern for narwhals given that changes to their endemic habitat are occurring at a faster rate than in other regions (Louis *et al*., 2020).

Here we analyse data collected by 8 animal-borne sound and movement tags deployed in East Greenland to model the sound production rates of narwhals. The reported acoustic cue rates for narwhals will contribute to improved density estimates as derived from passive acoustic data.

### II. MATERIALS AND METHODS

#### A. Field tagging

Narwhals were live-captured each August in 2013-2016, and 2019, at the Hjørnedal field station, located near the southwestern tip of Milne Land in the Scoresby Sound fjord complex (Figure 1), see (Blackwell *et al*., 2018). Tagging with sound and movement biologging tags (Acousonde*^T M^*, Acoustimetrics, California, USA) occurred near shore with set nets and the help of local Inuit hunters (Blackwell *et al*., 2018). Whale sex was determined by the presence (male) or absence (female) of a tusk as in Heide-Jørgensen *et al*. (2015). Six females and two males were tagged with Acousondes using suction cups, on the rear dorsal side (Figure 1), next to the dorsal ridge. Tags were further secured with a line and corrodible magnesium link which released the tag after *∼*4-9 days (see Blackwell *et al*. (2018)). For easier reference, each tagged animal was given a name (Table I): Freya, Thora, Mára, Frida, Balder, Eistla, Mutti, and Jonas.

**FIG. 1.**
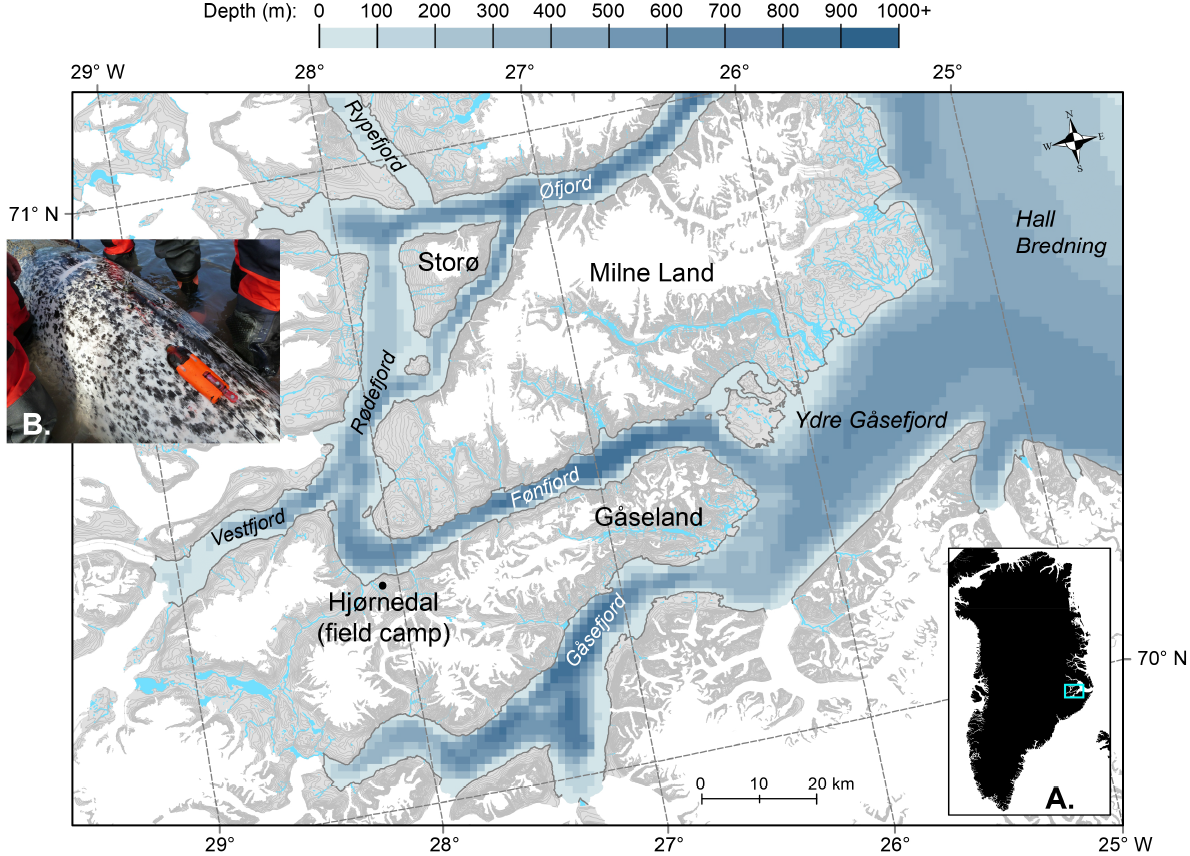
Map of study area, showing the fjords and inlets of Scoresby Sound, East Greenland (see inset (A)). Inset (B) shows the placement of the Acousonde tag on the female Freya.

**TABLE I.** Summary table with information on the tagged animals and their records. The variable *time* is the total number of days of each tag record. The whales’ silent period (in days), between release from tagging and the onset of echolocation-click’s production, is represented by *t_f_* . (These data were discarded from the analysis.) The actual time used in the analysis is given by *t_a_* in each record, and *samples* are the number of one-second samples manually screened for clicks.

| Narwhal ID | sex | year | time | $t_f$ | $t_a$ | samples | acoustic sample rate (Hz) |
| --- | --- | --- | --- | --- | --- | --- | --- |
| Freya | Female | 2013 | 2.84 | 1.53 | 1.31 | 200 | 25811 /154868 |
| Thora | Female | 2014 | 4.23 | 0.39 | 3.83 | 150 | 25811 /154868 |
| Mára | Female | 2014 | 0.42 | 0.37 | 0.05 | 10 | 25811 /154868 |
| Frida | Female | 2015 | 3.44 | 1.12 | 2.32 | 149 | 25811 |
| Eistla | Female | 2016 | 4.21 | 0.75 | 3.46 | 150 | 25811 /154868 |
| Balder | Male | 2016 | 8.71 | 2.61 | 4.35 | 194 | 25811 |
| Jonas | Male | 2019 | 9.34 | 0.28 | 6.10 | 200 | 25811 |
| Mutti | Female | 2019 | 6.38 | 1.21 | 3.02 | 150 | 25811 |
| Total | - | - | 34.46 | 8.26 | 24.44 | 1203 | - |

#### B. Permitting

Permission for capturing, handling, and tagging of narwhals was provided by the Government of Greenland (Case ID 2010–035453, document number 429 926). The project was reviewed and approved by the IACUC of the University of Copenhagen (17 June 2015). Access and permits to use land facilities in Scoresby Sound were provided by the Government of Greenland. No protected species were sampled. The secondary analysis of these data (as previously published in Blackwell *et al*. (2018) and Tervo *et al*. (2021) was approved by the University of St Andrews’ School of Biology Ethics Committee (BL16585).

#### C. Sampling scheme

Acoustic recordings were either continuous (4 whales, 25811 Hz) or duty-cycled (4 whales), alternating between low and high sampling rates (8 min at 25811 Hz, 7 min at 154868 Hz, see Table I and Blackwell *et al*. (2018)). Depth was sampled at 10 Hz in all tags, and then averaged to a 1 Hz sampling rate. Thora, Eistla, Jonas, and Balder’s tags detached after memory capacity was reached, while Freya, Frida, and Mutti’s tags detached from the whale before recording ended; Mára’s record ended prematurely due to a power reset. Jonas record lasts *∼*3 days longer but at the time of the analyses presented here those data had not yet been processed for presence / absence of clicks.

#### D. Description of tag data

After tagging, narwhals have been observed to remain silent for a variable amount of time, presumably as a consequence of the handling procedure (Blackwell *et al*., 2018). To avoid including such periods which would underestimate cue rates (e.g. Shuert *et al*., 2021), we excluded from the analyses any data preceding the start of echolocation, corresponding to between 0.39 and 2.61 days per tag deployment (Table I). Our assumption was that the tagged narwhal was foraging as normal when echolocation was observed to have resumed, and tag-on effects were diminished. The tag-on effects have recently been evaluated by Nielsen *et al*. (2023) and Shuert *et al*. (2021).

Prior to the work described in this manuscript, Blackwell *et al*. (2018) annotated each acoustic record for periods of echolocation (regular clicks), generally starting during the descent of foraging dives and ending during the ascent. Within these clicking bouts, silent periods had to exceed 10 seconds in duration to be labelled as “non-clicking” (Blackwell *et al*., 2018). In a given second from a clicking period, the whale may produce one or multiple echolocation clicks, or no clicks at all. Hence the original dataset, shared by Blackwell *et al*. (2018), consisted of time-depth profiles, at one-second resolution, with an associated time, clicking state (obtained by the method previously explained, where 1 = clicking, 0 = non-clicking), and depth for each second (Figure 2). To obtain an echolocation click production rate from these data we must first estimate the mean number of clicks produced per second during clicking periods.

**FIG. 2.**
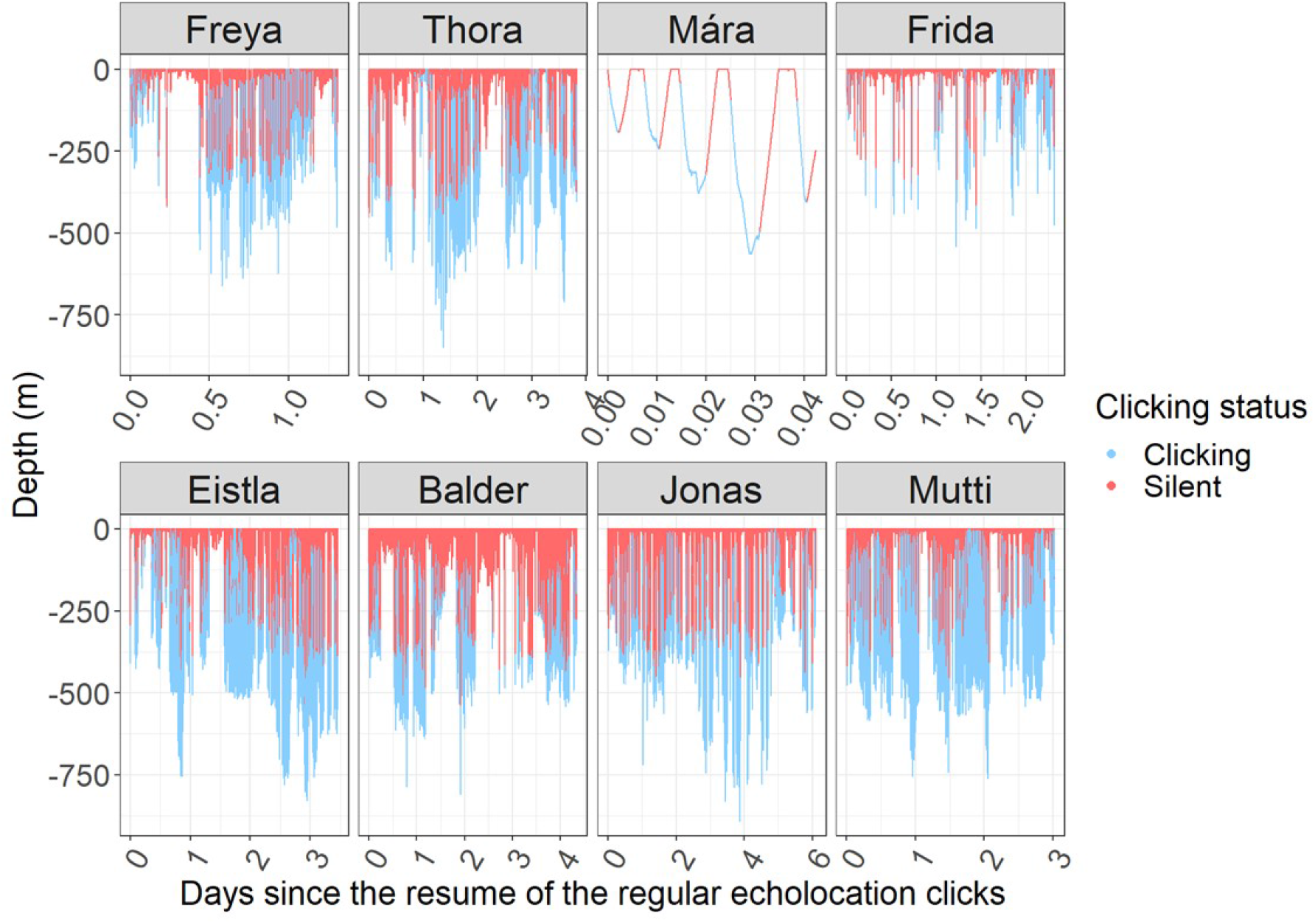
Depth profiles for each narwhal. Each second is coloured depending on it being in a clicking period (in blue) or in a non-clicking period (in pink).

#### E. Sample selection during clicking periods

The first step to obtain an overall cue rate is to determine the average clicking rate during clicking periods. This was done through systematic random sampling of all clicking seconds in each record, where the length of the record determined the number of samples taken, within four predefined categories, 10, 100, 150, or 200 samples (Table I). This method was chosen to avoid the burden of laborious human annotation of the full duration of the acoustic records. The systematic random sampling has a random start, so to evaluate the potential variability associated with a different random start, Freya’s dataset was sampled twice (100 samples each time). The comparison of the results obtained by Freya’s two datasets did not show significant differences and showed the sample size to be sufficient (for more information see the results in the Supplementary Material). Therefore, Freya was then used as a benchmark for the sampling for the other narwhals that had different recording durations.

#### F. Click counting method

Echolocation clicks in each sampled second were counted using sound pressure time series of the collected data in MT Viewer (a custom-written program for analysis of Acousonde data) (Burgess *et al*., 1998). To allow for a clearer view of the clicks and reduce the amount of flow noise, the acoustic data collected during low-frequency sampling were first high-pass filtered at 1.5 kHz (Blackwell *et al*., 2018). This high-pass filtration step during analysis was unnecessary for the acoustic data sampled at high frequency, for which the Acousonde has a built-in high-pass filter at 9.85 kHz (Blackwell *et al*., 2018).

Each second evaluated for clicks is referred to as a sampling unit. Only sampling units exclusively containing regular echolocation clicks were used for further cue rate analysis. When narwhals transition from regular clicks to feeding buzzes, there is a progressive decrease in the recorded click amplitude and an increase in the click repetition rate, during which click rates can exceed 300 clicks/s (Rasmussen *et al*., 2015). The reduced source level of odontocete buzz clicks renders them harder to detect on non-animal-borne PAM sensors. For this reason, samples occurring during buzzes were recorded as having zero clicks, starting once click density exceeded 20/s. An indicator variable was recorded to allow distinction between true zeros (absence of clicks) and buzz-induced zeros.

If a selected sample time was unavailable (e.g., Acousonde transitioning between low- and high-frequency recording, which takes 3-4 seconds), an adjacent sampling unit was selected instead. We assume these seconds are missing at random with respect to the echolocation click production rate and hence do not induce bias in reported estimates. In addition, in a handful of one-second samples (1 for Frida, 6 for Balder) clicks could not be counted in the samples because of poor signal-to-noise ratios — these samples were removed, resulting in lower sample sizes for some whales (Table I). Again, we assume these would be missing at random with respect to click production.

#### G. Estimation of echolocation click rate

Assuming the available tagged animals correspond to a random sample of narwhals in the population, the click production, *r*, can be estimated using the average of click production rates for each tag record, that is

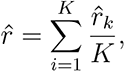

where *r̂_k_* is the estimated cue rate for tag record *k*, itself estimated as

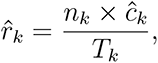

where *n_k_*represents the number of seconds initially recorded as clicking periods from tag *k*, *ĉ_k_* is the mean number of clicks per second in sampling data from tag *k* and *T_k_* is the total number of seconds of the acoustic recording considered in tag *k*.

In our case, where some tag records are much longer than others, it might be more sensible to consider a weighted average, weighted by tag record duration. The implications of using weighted vs. unweighted averages in this context are discussed in Marques *et al*. (Submitted) (see also the discussion section). The estimate for cue rate here becomes:

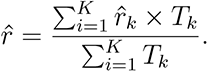

To evaluate how precision at the tag deployment level increases with the number of samples selected we plotted the coefficient of variation (CV) of the cue rate estimates for each individual tag record as a function of sample size, using a bootstrap procedure. For each sample size we computed the cue rate and corresponding CV based on 1000 resamples with replacement.

We used Hierarchical Generalized Additive Models (GAM) (Wood, 2011) to model, at second-by-second resolution, both 1) the probability of being in a clicking state and 2) the number of clicks, as a function of both time since the onset of foraging (i.e., the effective start of the record in this analysis) and depth. The first model considered a Binomial response and logit link function, while the second model considered a Poisson response and log link function. For both models we considered either a common smoothing or a separate smoothing by tag (Pedersen *et al*., 2019). Model selection was based on AIC.

Data analysis was implemented in software R (Team, 2021). GAM models were implemented using function gam, from the package mgcv (Wood, 2011). The code and data required to reproduce all the results in this paper are shared in the github repository X.

## III. RESULTS

From the preliminary analysis done using data containing information about the clicking phases, depth on second-by-second resolution we obtained the following results:

A total of 24.44 days of on-whale acoustic recordings were available after whales resumed their regular echolocation clicks. At that point whales performed deep dives interspersed with periods at the surface, and spent on average 32% of their time (se=0.09, cv=0.29) clicking, ranging from 19% for Frida to 47% for Mutti.

From the analysis done using the sample data that contain information about the number of clicks per second inside the clicking period, we obtained the following results:

The CV on the mean number of clicks decreased with the number of samples selected, reaching values lower than 0.2 at less than 25 one-second-duration random samples, and with values lower than 0.1 at the maximum number of samples (Figure 3). This happened for all narwhals except for Mára, who only had 10 samples available given the short tag duration.

**FIG. 3.**
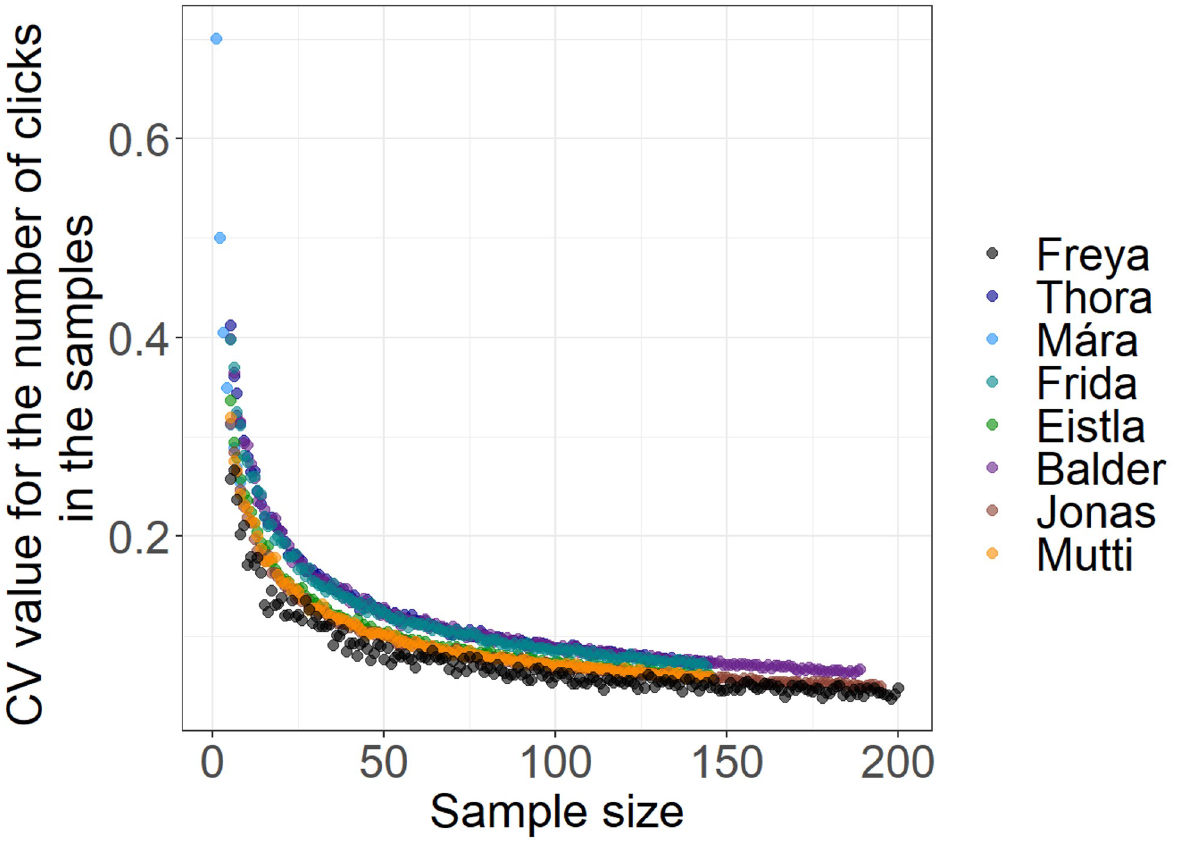
CV of the mean number of clicks per second as a function of the number of samples considered, for each tagged whale.

We consider for further inference and interpretation the GAM models with the lowest AIC (Table II). The first one was the model used to predict the proportion of time spent in a clicking state, with depth and narwhal ID being included in the model (AIC = 83993.21). The probability of being in a clicking state increases with depth, to near certain (P = 1) at depths below about 500 meters (Figure 4). The second model predicted the number of clicks per second as a function of depth, days since the first onset of regular echolocation clicks and narwhal ID being included in the model (AIC = 6732.86). The clicking rate (clicks / second) shows a slight and non-significant decreasing pattern over tag deployment time and an increasing pattern with depth, with considerable variation across animals being strongly supported by the most parsimonious model (Figure 5). The model results show that time since the first onset of regular echolocation clicks does not seem to influence most of the narwhals, the only ones that show significant differences are Balder, Freya and Frida. The model also predicts that depth influences the production of clicks for all the narwhals except for Mára.

**FIG. 4.**
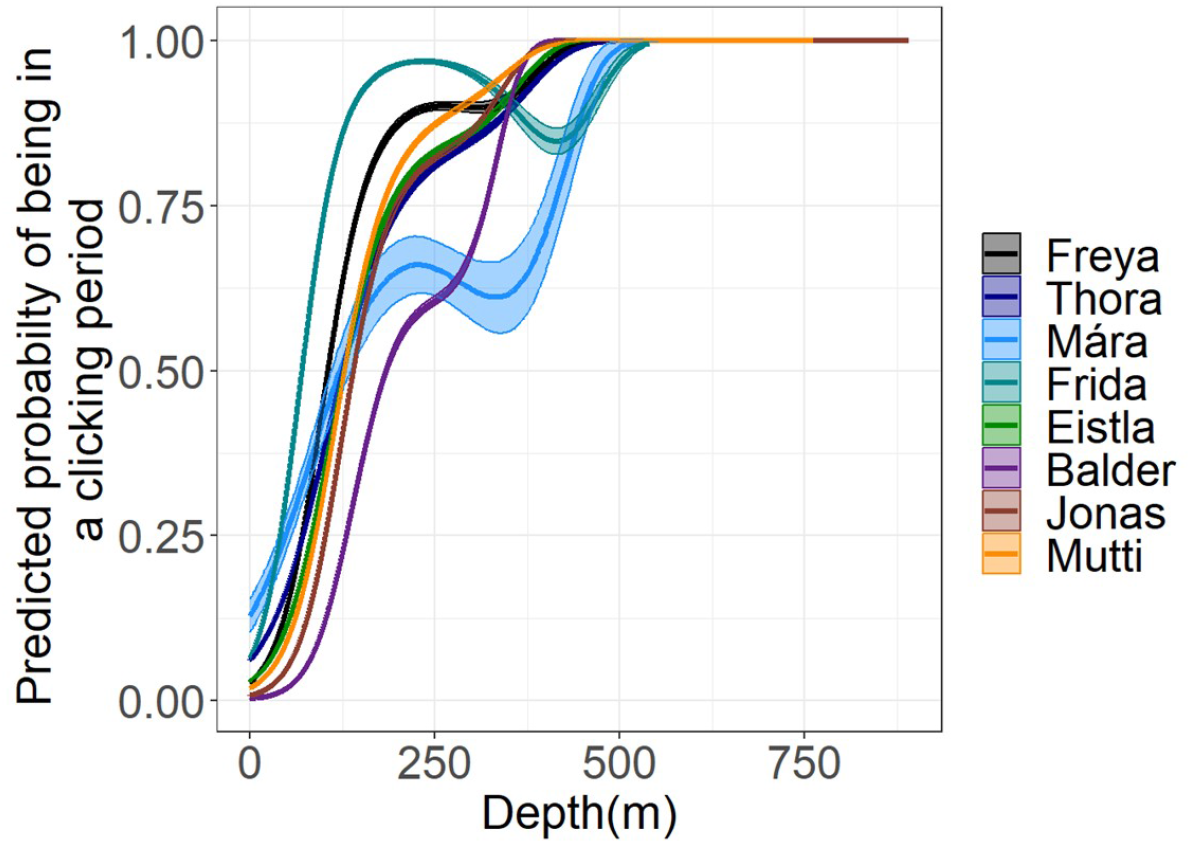
The whale-specific estimated probability of being in a clicking state as a function of depth. Lines show predicted values and the shaded areas correspond to 95% confidence intervals.

**FIG. 5.**
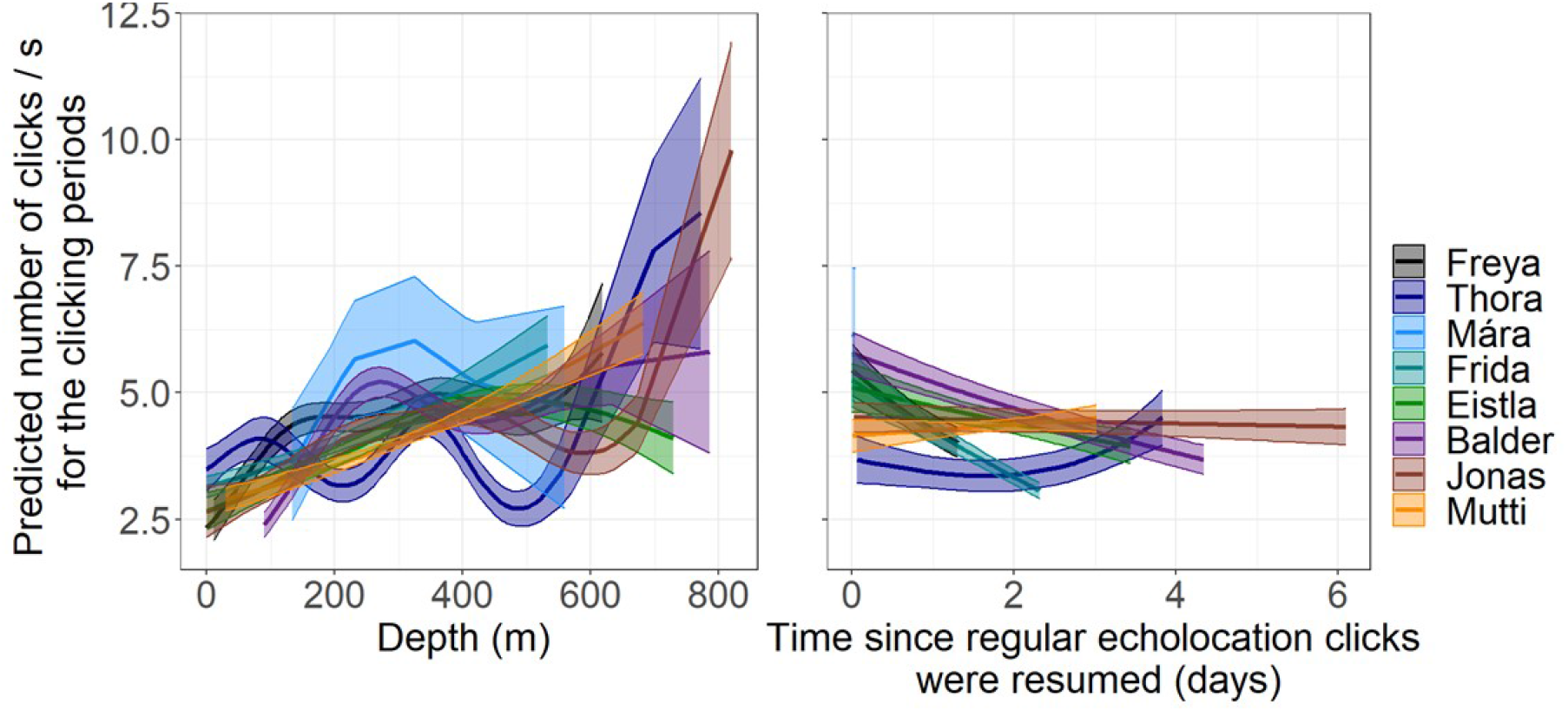
Clicking rate (clicks / s) as a function of days since initial onset of regular echolocation clicks (right) and depth (left), while in a clicking state. Lines show predicted values and shaded areas correspond to 95% confidence intervals.

**TABLE II.** Summary table with information on the different models and their AIC. *Response variable* is the variable being modeled and *Explanatory variables* corresponds to the variables used in the model, where depth is in meters and the time since initial onset of regular echolocation clicks, in seconds.

| Response variable | Explanatory variables | AIC |
| --- | --- | --- |
| Clicking period (1)/ Silent period (0) | depth by narwhal ID | 83993.21 |
| Clicking period (1)/ Silent period (0) | depth | 91391.09 |
| Number of clicks per second | depth and time by narwhal ID | 6732.86 |
| Number of clicks per second | depth by narwhal ID | 6771.62 |
| Number of clicks per second | depth | 6787.72 |
| Number of clicks per second | depth and time | 6807.83 |

The predicted number of clicks for the clicking periods is overlaid on the narwhals’ depth profiles in Figure 6.

**FIG. 6.**
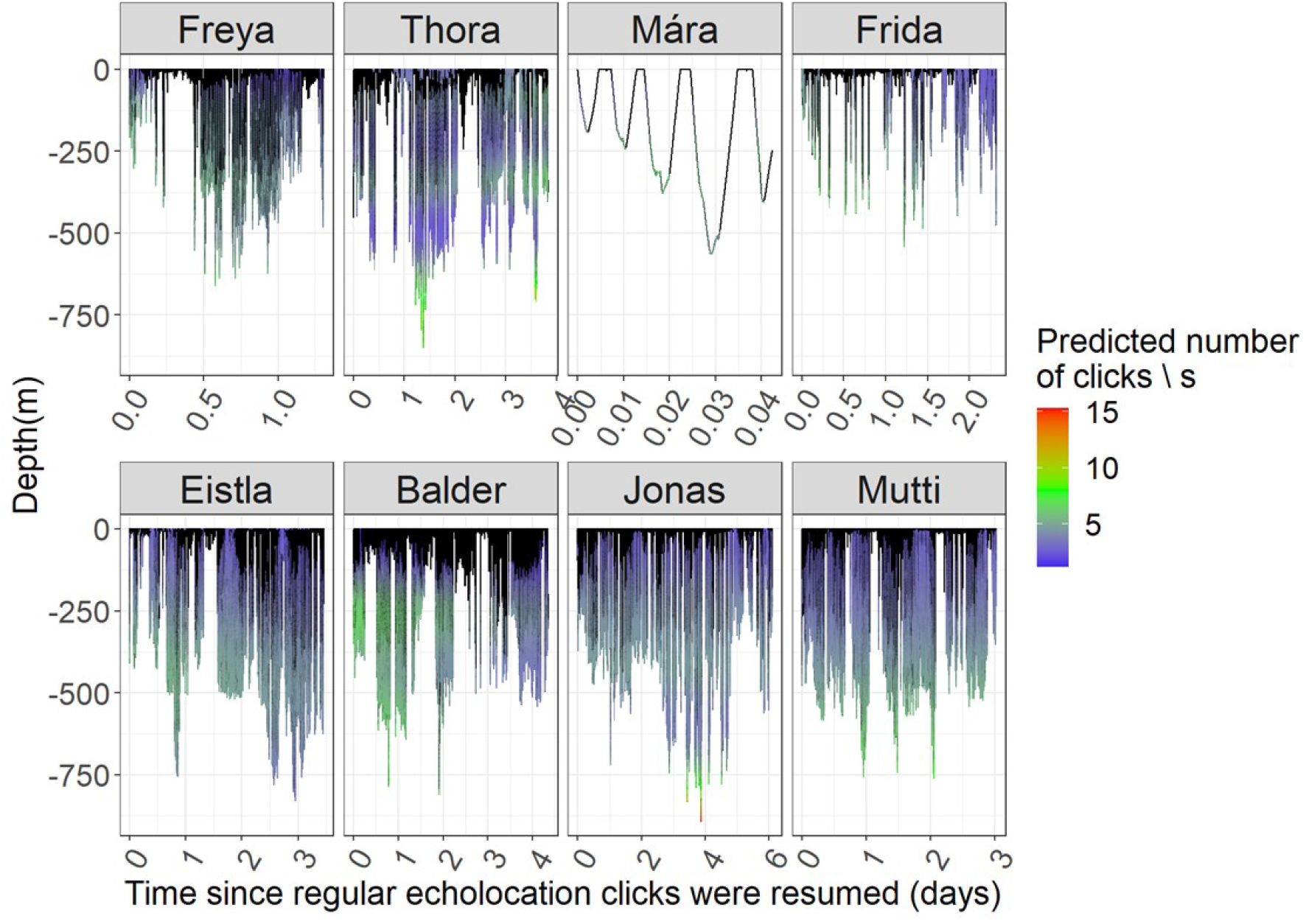
Depth profiles with colour proportional to the predicted number of clicks during the clicking periods. Black corresponds to silent periods.

Individual estimated click production rates range from 0.72 clicks / second, for Frida, to 1.98 clicks / second for Mutti, disregarding the value 2.12 for Mára, based on only 10 samples (Table III). Overall, the echolocation click production rate, considering the weighted average, was estimated to be 1.28 clicks per second (95% CI = 0.98, 1.58). For comparison, the standard average was 1.40 clicks per second (95%CI = 1.01, 1.80).

**TABLE III.**
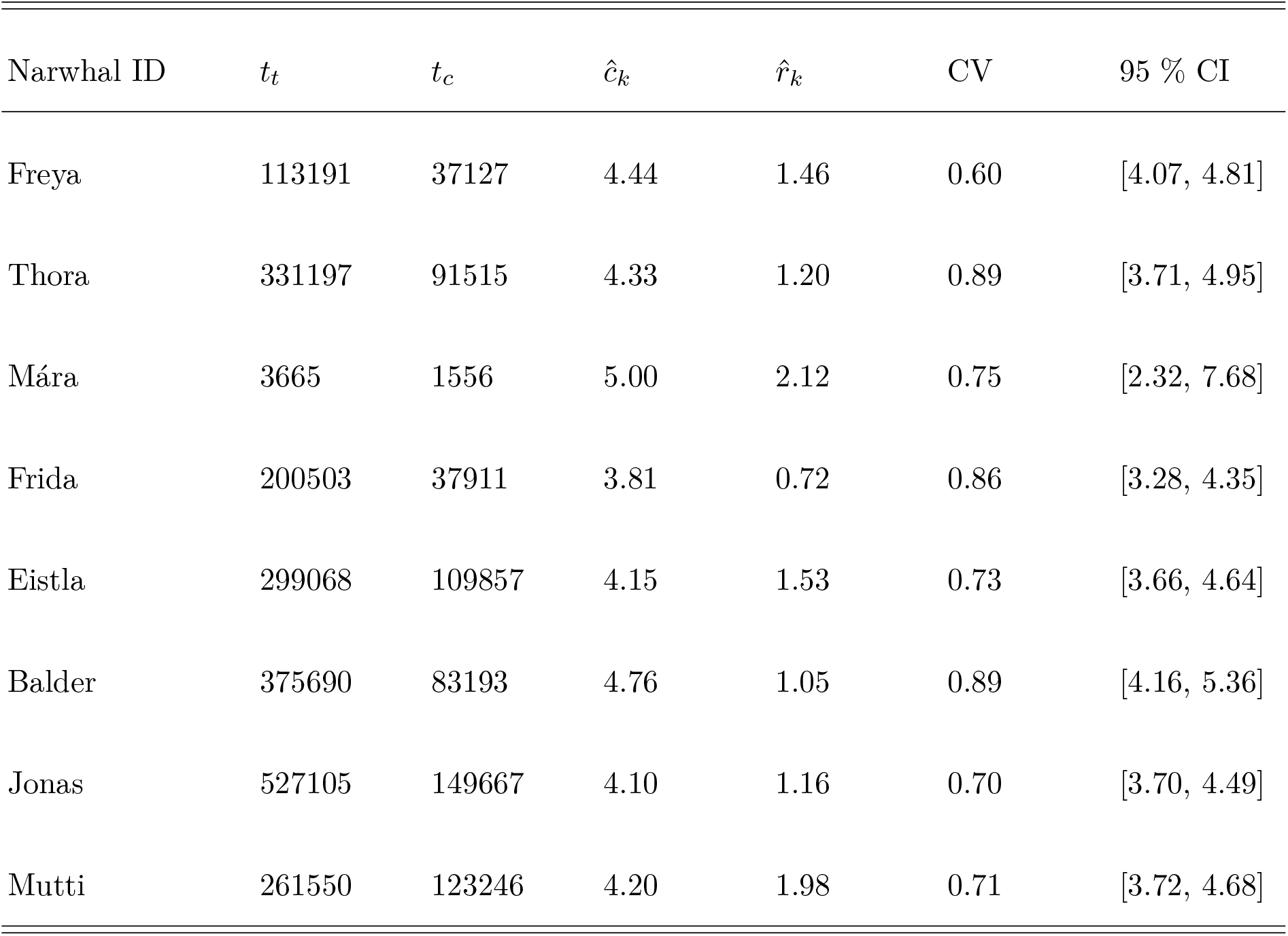
Cue rates (*r̂_k_*) obtained for each tagged whale. *N arwhal ID* is followed by tag record duration (*t_t_*, in seconds), the total time spent clicking(*t_c_*, in seconds) and the mean number of clicks produced per second, *ĉ_k_*. Also shown the per animal coefficient of variation (*CV*) and 95% confidence intervals (*CI*) are.

## IV. DISCUSSION

Our main objectives were 1) to provide a first cue rate for narwhals to inform passive acoustic cue counting estimates via cue counting and 2) to understand how cue rate might vary over time and with depth. Given the structure of the available data, we did the latter using a two-stage approach, first modelling the probability of being in a clicking state, and then, conditional on being in said state, modelling the number of clicks per second.

Our cue rate estimates were consistent, irrespective of whether the average was weighted (based on usable tag record duration), or not, with largely overlapping 95% confidence intervals. This is reassuring since it means that the reported results were not influenced by a somewhat arbitrary decision of whether or not to weigh the averages by tag record duration. Weighting (vs. non weighting) might be optimal depending on whether one can safely assume that longer tag records provide more reliable information about the average cue rate than shorter tag records. However, despite a seemingly easy question on the surface, this becomes a potentially complicated problem in practice. A longer recording should provide more reliable data on a given animal, but there is a limit to such information gain. An absurd example is useful to illustrate our point. In the limit, if one weighs for tag duration, and one has 100 tags with 1 hour each and 1 tag with a year, the weighted average would be almost solely driven by the longer tag. This would certainly be undesirable, highlighting that what is at stake is how variable animals are over time compared to the variability between animals. Marques *et al*. (Submitted) further discuss different approaches to estimating cue rates from tag data. In our case, Mára’s tag only contained reliable information for 1 hour and never reached depths below 600 meters. Therefore, her results are less reliable than her counterparts’. For this reason, for narwhal PAM density estimates that may require a cue rate estimate, we suggest the following weighted average cue rate estimate and corresponding precision: 1.28 clicks/second (standard error: 0.13; CV: 0.10; 95% CI [0.98,1.58]).

The increasing probability of being in a clicking state with depth, as well as the number of clicks while clicking also increasing with depth, were expected, given that narwhals are considered to be deep divers, with echolocation bouts and presumed foraging occurring at depth (Blackwell *et al*., 2018). Having these results, it would be interesting to compare them with another model that uses the depth of the seabed (which can be assessed using the GPS position for the narwhals, accessible by Argos transmitter (Blackwell *et al*., 2018)) instead of diving depth, allowing us to put clicking into further ecological context, but this would require analysis beyond the scope of this paper.

Frida, Freya, and Balder’s data suggest a negative linear tendency in the number of clicks with time (Figure 5). This unexpected result may be explained by the fact that Freya’s and Frida’s records were comparatively short due to the early detachment of their tags (Table I). After tagging, Balder took more than 2.58 days to begin foraging, which indicates that it may have taken him longer to recover from the tagging process. For these reasons, we avoid over-interpreting the data. Our results, with a consistent cue rate obtained with only 8 tag records, indicate that echolocation clicks can probably be suited as ideal cues for density estimates by PAM for narwhals. However, this dataset also highlights how long-term tags might be fundamental to understand cue rates, and in particular to tease apart relevant unanticipated tag-on effects.

A preferable approach to the laborious manual annotation of the acoustic records would be to use a reliable automated click detector, which would yield the times of every regular click emitted by the whales. We are therefore currently developing an automated procedure for narwhal click detection.

This study was based on a small number of tag records, from a single location and time of the year. It would be desirable to investigate whether the values found here are stable enough to be used for other times and places for which a density estimate via PAM cue counting may be attempted for narwhals. For the time being, whether the cue rate reported here is adequate under different settings remains an untested assumption. Such cue rates from other times and places are often used when lacking better alternatives (see (e.g. Marques *et al*., 2011) for an example with right whales). Three narwhal populations—in East Greenland, West Greenland, and Northern Hudson Bay—show different dive behaviors as a function of season, have different diets, and use different water column depths for different purposes, which may result in differences between the sounds produced in distinct locations or even between individuals (Marcoux *et al*., 2012; Watt *et al*., 2015). Indeed, variations in the acoustic detection of narwhals have been observed between PAM recording locations, corresponding with different stages of their biannual north/south migration (Frouin-Mouy *et al*., 2017). Furthermore, annual variability of ambient noise levels within long-term PAM recording stations in the Arctic may influence both narwhal behaviour and acoustic detection capabilities (Ladegaard *et al*., 2021). It may be inappropriate to extrapolate our cue production rate to areas where narwhals have different depth preferences, hence, cue productions rate might have to be estimated separately for each population. All of the narwhals in this study were tagged in the summer. This might lead to a potential bias when extending this cue rate, here calculated exclusively from the month of August, into density estimates for other seasons. For example, Chambault *et al*. (2023) show that in data also exclusively obtained during summer months, narwhal’s prey capture efficiency was much lower than expected, leading them to presume that narwhals sustain their bioenergetic demands by foraging more at other times of the year. Reduced prey density, extreme prey selection and varied energetic needs at different times of year could all have contributed to a further nuanced use of the value presented here.

This study used tag data obtained following live captures and handling of animals. The whales’ acoustic behavior following release was unusually silent, and for that reason we removed up until the onset of regular echolocation clicks. In Shuert *et al*. (2021) and Nielsen *et al*. (2023) the behavioural responses of individuals following tagging and capture events were evaluated. In general, individual narwhals recovered quickly from these events (Nielsen *et al*., 2023; Shuert *et al*., 2021). On the other hand, significant inter-individual variability in the magnitude of the response was observed (Nielsen *et al*., 2023; Shuert *et al*., 2021). Most individuals appeared to recover in the first 8–9 hours following release (Nielsen *et al*., 2023; Shuert *et al*., 2021). However, both studies have shown that the handling time influences the post-release behaviour, where individuals held for more than 40 minutes (in Shuert *et al*. (2021)) or 58 minutes (in Nielsen *et al*. (2023)) exhibited the largest change in activity following the handling events. Those results may differ somewhat from the ones presented here, since in this study we assume the recovery of the animal is given by the first onset of regular echolocation clicks, a conservative approach, while Shuert *et al*. (2021) focused on accelerometry-derived behavioural metrics of activity levels, energy expenditure, and swimming behaviour to obtain this information and Nielsen *et al*. (2023) used quantile regression and relative entropy.

Preliminary unreported models showed no effect of sex or animal length on cue rates, but the power to find existing differences was unsurprisingly low, so we abstained from presenting those results here. It is important to know to what extent cue production rates are potentially influenced by animal-dependent factors like size, sex, or age, which could cast doubts about the reliability of our estimate given the small sample size and the increased likelihood of finding animals (as per Webster and Rutz (2020)). Care should be taken also with respect to non-individual animal external factors, including prey distribution or prey abundance. As an example, Watt *et al*. (2013, 2015) report differences in diving behaviour across sites, linking them to dietary preferences. These will have direct consequences on whether and how the cue rates reported here can be generalized to other areas.

Narwhals occur in the Arctic which is hard to reach and where weather is often poor, showing elusive behaviour and spending much of their time submerged, over a wide area at low densities. For these reasons, traditional visual survey methods are difficult to carry out in their habitats (Marques *et al*., 2009). As a result, PAM methods may be promising and could be highly effective (Marques *et al*., 2009). Our results represent a first step to obtain a cue production rate estimate for narwhals, opening the door to PAM-mediated cue counting approaches to estimate their density. An accurate density estimate for narwhals, which is needed for their management and conservation (Podolskiy and Sugiyama, 2020), may be best obtained using PAM, which depends on the reliability of the cue rates if based on cue counting methods. Climate change has brought an increasing number of threats to these animals, such as an ice-free Arctic with a corresponding increase in shipping; therefore, the importance of monitoring the state of narwhal populations over the years should not be understated (Podolskiy and Sugiyama, 2020; Shuert *et al*., 2022). In addition to an increase in shipping, the ice-free Arctic will also have more room for oil exploration (Williams *et al*., 2022), further increasing human pressure on an already fragile system. To our knowledge, PAM has not been used for narwhal density estimation, and mean click rates presented here get us one step closer to obtaining a PAM density estimate, in particular for the Scoresby Sound sub-population. The missing stepping stone, however, is knowledge about the effective detection area of PAM sensors, or equivalently, the probability of acoustic detection of a narwhal click around a sensor. This could be obtained in several ways, including distance sampling (e.g. Marques *et al*., 2011), spatially explicit capture-recapture (e.g. Stevenson *et al*., 2021), with trial-based methods (e.g. Marques *et al*., 2009), or based on acoustic propagation methods (e.g. Küsel *et al*., 2011). Density estimates derived this way for animals with highly directional acoustic cues (e.g., odontocete echolocation clicks) would further benefit from information on biosonar parameter quantification and the echolocation beam pattern, which have recently been reported by Zahn *et al*. (2021) and Koblitz *et al*. (2016), respectively. We look forward to seeing studies that address these questions, and we hope the cue rate provided here might inspire researchers to sooner rather than later try to estimate narwhal abundance using PAM cue counting.

## ACKNOWLEDGMENTS

This research was conducted under the ACCURATE project, funded by the US Navy Living Marine Resources program (contract no. N3943019C2176). TAM and CSM thank partial support by CEAUL funded through the project ACCURATE4CEAUL. We also thank Rikke G. Hansen for her help with Figure 1.

## Notes

https://github.com/MCarolinaMarques/Narwhal-echolocation-click-rates-article-analysis.git

### Competing Interest Statement

The authors have declared no competing interest.

## REFERENCES

Blackwell, S. B., Tervo, O. M., Conrad, Hansen, R. G., and Ditlevsen (2018). “Spatial and temporal patterns of sound production in East Greenland narwhals,” PLoS One 13(6), e0198295, https://doi.org/10.1371/journal.pone.0198295.

Burgess, W. C., Tyack, P. L., Boeuf, B. J. L., and Costa, D. P. (1998). “A programmable acoustic recording tag and first results from free-ranging northern elephant seals,” Deep Sea Research Part II: Topical Studies in Oceanography 45(7), 1327–1351, https://doi.org/10.1016/S0967-0645(98)00032-0.

Chambault, P., Blackwell, S. B., and Heide-Jørgensen, M. P. (2023). “Extremely low seasonal prey capture efficiency in a deep-diving whale, the narwhal,” 19(2), https://doi.org/10.1098/rsbl.2022.0423.

Frouin-Mouy, H., Kowarski, K., and Martin (2017). “Seasonal trends in acoustic detection of marine mammals in Baffin Bay and Melville Bay, Northwest Greenland,” Arctic 59–76, https://www.jstor.org/stable/26379724.

Harris, D. V., Miksis-Olds, J. L., Vernon, J. A., and Thomas, L. (2018). “Fin whale density and distribution estimation using acoustic bearings derived from sparse arrays,” The Journal of the Acoustical Society of America 143(5), 2980–2993, https://doi.org/10.1121/1.5031111.

Heide-Jørgensen, M. P., Nielsen, N. H., Hansen, Blackwell, S. B., and Jørgensen, O. A. (2015). “The predictable narwhal: satellite tracking shows behavioural similarities between isolated subpopulations,” Journal of Zoology 297(1), 54–65, https://doi.org/10.1111/jzo.12257.

Johnson, M. P., and Tyack, P. L. (2003). “A digital acoustic recording tag for measuring the response of wild marine mammals to sound,” IEEE journal of oceanic engineering 28(1), 3–12, https://doi.org/10.1109/JOE.2002.808212.

Koblitz, J. C., Stilz, P., Rasmussen, M. H., and Laidre, K. L. (2016). “Highly Directional Sonar Beam of Narwhals (*Monodon monoceros*) Measured with a Vertical 16 Hydrophone Array,” PLOS ONE 11(11), e0162069, https://doi.org/10.1371/journal.pone.0162069.

Küsel, E. T., Mellinger, D. K., Thomas, L., Marques, T. A., Moretti, D., and Ward, J. (2011). “Cetacean population density estimation from single fixed sensors using passive acoustics,” The Journal of the Acoustical Society of America 129(6), 3610–3622, https://doi.org/10.1121/1.3583504.

Ladegaard, M., Macaulay, J., Simon, M., Laidre, K. L., Mitseva, A., Videsen, S., Pedersen, and Madsen, P. T. (2021). “Soundscape and ambient noise levels of the Arctic waters around Greenland,” Scientific Reports 11(1), 23360, https://doi.org/10.1038/s41598-021-02255-6.

Laidre, K. L., Stirling, I., Lowry, L. F., Wiig, Ø., Heide-Jørgensen, M. P., and Ferguson, S. H. (2008). “Quantifying the sensitivity of Arctic marine mammals to climate-induced habitat change,” Ecological Applications 18(sp2), S97–S125, https://doi.org/10.1890/06-0546.1.

Louis, M., Skovrind, M., Castruita, J. A. S., Garilao, C., Kaschner, K., Gopalakrishnan, S., Haile, J. S., Lydersen, C., Kovacs, K. M., Garde, E., Heide-Jørgensen, M. P., Postma, L., Ferguson, S. H., Willerslev, E., and Lorenzen, E. D. (2020). “Influence of past climate change on phylogeography and demographic history of narwhals, *Monodon monoceros*,” Proceedings of the Royal Society B: Biological Sciences 287(1925), 20192964, https://doi.org/10.1098/rspb.2019.2964.

Macaulay, J. D., Malinka, C. E., Gillespie, D., and Madsen, P. T. (2020). “ High resolution three-dimensional beam radiation pattern of harbour porpoise clicks with implications for passive acoustic monitoring,” The Journal of the Acoustical Society of America 147(6), 4175–4188, https://doi.org/10.1121/10.0001376.

Marcoux, M., Auger-Méthé, M., and Humphries, M. M. (2012). “Variability and context specificity of narwhal (*Monodon monoceros*) whistles and pulsed calls,” Marine Mammal Science 28(4), 649–665, https://doi.org/10.1111/j.1748-7692.2011.00514.x.

Marques, T. A., Marques, C. S., and Gkikopoulou, K. C. (Submitted). “A cautionary tale about estimating acoustic cue rates for deep divers: a sperm whale example,” Submitted to Journal of the Acoustical Society of America.

Marques, T. A., Munger, L., Thomas, L., Wiggins, S., and Hildebrand, J. A. (2011). “Estimating North Pacific right whale (*Eubalaena japonica*) density using passive acoustic cue counting,” Endangered Species Research 13, 163–172, https://doi.org/10.3354/esr00325.

Marques, T. A., Thomas, L., Martin, S. W., Mellinger, D. K., Ward, J. A., Moretti, D. J., Harris, D., and Tyack, P. L. (2013). “Estimating animal population density using passive acoustics,” Biological reviews 88(2), 287–309, https://doi.org/10.1111/brv.12001.

Marques, T. A., Thomas, L., Ward, J., DiMarzio, N., and Tyack, P. L. (2009). “Estimating cetacean population density using fixed passive acoustic sensors: An example with Blainville’s beaked whales,” The Journal of the Acoustical Society of America 125(4), 1982–1994, https://doi.org/10.1121/1.3089590.

Nielsen, L. R., Tervo, O. M., Blackwell, S. B., Heide-Jørgensen, M. P., and Ditlevsen, S. (2023). “ Using quantile regression and relative entropy toassess the period of anomalous behaviour ofmarine mammals following tagging,” Ecology and Evolution 13(4), e9967, https://doi.org/10.1002/ece3.9967.

Nowacek, D. P., Christiansen, F., Bejder, L., Goldbogen, J. A., and Friedlaender, A. S. (2016). “Studying cetacean behaviour: new technological approaches and conservation applications,” Animal Behaviour 120, 235–244, https://doi.org/10.1016/j.anbehav.2016.07.019.

Pedersen, E. J., Miller, D. L., Simpson, G. L., and Ross, N. (2019). “Hierarchical generalized additive models in ecology: an introduction with *mgcv*,” PeerJ 7, e6876, https://doi.org/10.7717/peerj.6876.

Podolskiy, E. A., and Sugiyama, S. (2020). “Soundscape of a Narwhal Summering Ground in a Glacier Fjord (Inglefield Bredning, Greenland),” Journal of Geophysical Research: Oceans 125(5), https://doi.org/10.1029/2020JC016116.

Rasmussen, M. H., Koblitz, J. C., and Laidre, K. L. (2015). “Buzzes and High-Frequency Clicks Recorded from Narwhals (*Monodon monoceros*) at Their Wintering Ground,” Aquatic Mammals 41(3), 256–264, https://doi.org/10.1578/AM.41.3.2015.256.

Scheidat, M., Geelhoed, S. C. V., and Noort, C. (2019). “Feasibility of a Passive Acoustic Monitoring (PAM) network for Harbour Porpoises on the Dutch Continental Shelf,” Technical Report (Wageningen Marine Research), https://doi.org/10.18174/504076.

Shuert, C. R., Marcoux, M., Hussey, Dietz, R., and Auger-Méthé, M. (2022). “Decadal migration phenology of a long-lived Arctic icon keeps pace with climate change,” Proceedings of the National Academy of Sciences 119(45), e2121092119, https://doi.org/10.1073/pnas.2121092119.

Shuert, C. R., Marcoux, M., Hussey, N. E., Watt, C. A., and Auger-Méthé, M. (2021). “Assessing the post-release effects of capture, handling and placement of satellite telemetry devices on narwhal (*Monodon monoceros*) movement behaviour,” Conservation Physiology 9(1), https://doi.org/10.1093/conphys/coaa128.

Stevenson, B. C., van Dam-Bates, P., Young, C. K., and Measey, J. (2021). “A spatial capture–recapture model to estimate call rate and population density from passive acoustic surveys,” Methods in Ecology and Evolution 12(3), 432–442, https://doi.org/10.1111/2041-210X.13522.

Stocker, T. F., Qin, D., Plattner, G.-K., Tignor, M. M. B., Allen, S. K., Boschung, J., Nauels, A., Xia, Y., Bex, V., and Midgley, P. M. (2014). “Climate Change 2013: The physical science basis. contribution of working group I to the fifth assessment report of IPCC the inter- governmental panel on climate change,” https://doi.org/10.1017/CBO9781107415324.

Team, R. C. (2021). “R: A language and environment for statistical computing,” .

Tervo, O. M., Ditlevsen, S., Ngô, M. C., Nielsen, N. H., Blackwell, S. B., Williams, T. M., and Heide-Jørgensen, M. P. (2021). “Hunting by the Stroke: How Foraging Drives Diving Behavior and Locomotion of East-Greenland Narwhals (*Monodon monoceros*),” Frontiers in Marine Science 7, https://doi.org/10.3389/fmars.2020.596469.

Thomas, L., and Marques, T. A. (2012). “Passive Acoustic Monitoring for Estimating Animal Density,” Acoustics Today 8(3), 35, https://doi.org/10.1121/1.4753915.

Warren, V. E., Marques, T. A., Harris, D., Thomas, L., Tyack, P. L., de Soto, N. A., Hickmott, L. S., and Johnson, M. P. (2017). “Spatio-temporal variation in click production rates of beaked whales: Implications for passive acoustic density estimation,” The Journal of the Acoustical Society of America 141(3), 1962–1974, https://doi.org/10.1121/1.4978439.

Watt, C. A., Heide-Jørgensen, M. P., and Ferguson, S. H. (2013). “How adaptable are narwhal? A comparison of foraging patterns among the world’s three narwhal populations,” Ecosphere 4(6), 1–15, https://doi.org/10.1890/ES13-00137.1.

Watt, C. A., Orr, J. R., Heide-Jørgensen, M. P., Nielsen, N. H., and Ferguson, S. H. (2015). “Differences in dive behaviour among the world’s three narwhal *Monodon monoceros* populations correspond with dietary differences,” Marine Ecology Progress Series 525, 273–285, https://doi.org/10.3354/meps11202.

Webster, M. M., and Rutz, C. (2020). “How STRANGE are your study animals?,” Nature 582(7812), 337–340, https://doi.org/10.1038/d41586-020-01751-5.

Weilgart, L. (2017). “Din of the Deep: Noise in the Ocean and Its Impacts on Cetaceans,” in Marine Mammal Welfare (Springer International Publishing), pp. 111–124, https://doi.org/10.1007/978-3-319-46994-2_7.

Westbury, M. V., Petersen, B., Garde, E., Heide-Jørgensen, M. P., and Lorenzen, E. D. (2019). “Narwhal Genome Reveals Long-Term Low Genetic Diversity despite Current Large Abundance Size,” iScience 15, 592–599, https://doi.org/10.1016/j.isci.2019.03.023.

Williams, T. M., Blackwell, S. B., Tervo, O., Garde, E., Sinding, M.-H. S., Richter, B., and Heide-Jørgensen, M. P. (2022). “Physiological responses of narwhals to anthropogenic noise: A case study with seismic airguns and vessel traffic in the Arctic,” Functional Ecology 36(9), 2251–2266, https://doi.org/10.1111/1365-2435.14119.

Williams, T. M., Noren, S. R., and Glenn, M. (2011). “Extreme physiological adaptations as predictors of climate-change sensitivity in the narwhal, *Monodon monoceros*,” Marine Mammal Science 27(2), 334–349, https://doi.org/10.1111/j.1748-7692.2010.00408.x.

Wood, S. N. (2011). “Fast stable restricted maximum likelihood and marginal likelihood estimation of semiparametric generalized linear models,” Journal of the Royal Statistical Society: Series B (Statistical Methodology) 73(1), 3–36, https://doi.org/10.1111/j.1467-9868.2010.00749.x.

Zahn, M. J., Rankin, S., McCullough, J. L., Koblitz, J. C., Archer, F., Rasmussen, M. H., and Laidre, K. L. (2021). “Acoustic differentiation and classification of wild belugas and narwhals using echolocation clicks,” Scientific Reports 11(1), 22141, https://doi.org/10.1038/s41598-021-01441-w.

